# Embedding Perfusable Microchannel Networks in Photoclickable Bioresins via High-Resolution Digital Light Processing

**DOI:** 10.1101/2025.10.29.685256

**Authors:** Riccardo Rizzo, Thibault Sampon, Jackson K. Wilt, Paul P. Stankey, Jennifer A. Lewis

## Abstract

Light-mediated 3D bioprinting methods hold great promise for the generation of biomimetic microvasculature networks for applications ranging from organ-on-chip models to vascularized tissue constructs. While printing microvascular channels (≤100 µm in diameter) within large hydrogel volumes (≥1 cm³) is theoretically feasible, progress remains limited by the lack of suitable biocompatible photoresins. Here, we report the development of an optimized photoresin based on fish gelatin and photoclick crosslinking chemistry for bioprinting perfusable, embedded microvascular networks via high-resolution digital light processing (DLP). Specifically, our biocompatible matrix leverages the fast kinetics and negligible dark curing of thiol-norbornene crosslinking as well as the low viscosity and thermal stability of fish gelatin. Using pulsed illumination and a biocompatible radical scavenger (DMPO), we further minimize radical diffusion-induced blurring, enabling extended printing (>5 h). Finally, printing failures are reduced through the incorporation of a biocompatible surfactant (Poloxamer-188). Together, these advances open new avenues for printing perfusable biomimetic microvascular networks embedded in biocompatible hydrogel matrices.

## 1. Introduction

Human microphysiological systems (also known as organ-on-chip models) offer a promising *in vitro* platform for drug screening, as highlighted by the FDA Modernization Act 2.0.^1^ However, most organ-on-chip models are made from rigid materials assembled via soft lithography, which contain perfusable microchannels separated by porous membranes that support cell monolayers. The ability to pattern complex 3D microvascular networks at high resolution (≤100 µm channel diameter) in biocompatible hydrogels across cm-scale devices would usher in next generation models that better emulate organ-specific microphysiology.

Multiple materials deposition^2–7^ and light-based^8–12^ methods have been developed to address this challenge. In particular, light-based 3D printing is a promising approach due to its precise spatiotemporal control over photoactive material crosslinking.^13, 14^ While two-photon stereolithography (2P-SL) offers the highest resolution (≤1 µm),^10, 15–17^ point-by-point laser scanning coupled with the small work distance associated with high-magnification objectives currently results in long printing times for limited build volumes (∼1 mm^3^). By contrast, tomographic volumetric additive manufacturing (TVAM) enables rapid printing (10’s second) of cm-size objects, but has limited spatial resolution (>250 µm negative features).^18, 19^ To overcome such limitations, hybrid approaches are emerging, which integrate the rapid printing of large perfusable models via TVAM with high resolution ablation of microcapillary-like structures via 2P-SL.^16^

Digital-light-processing (DLP) is an alternate approach that bridges the gap between TVAM and 2P-SL with respect to both print speed and spatial resolution. There has been significant progress in DLP patterning of hydrogels,^14^ including the identification of resins with optimal viscosity, biocompatible photoabsorbers (e.g., tartrazine,^8, 20^ chlorophyllin,^20^ Ponceau4R^21, 22^) and advances that enable multimaterial,^23–25^ high-cell density,^9^ and soft hydrogel printing.^22^ To date, most of these studies used commercial devices with limited printing resolution (i.e., 50-100 µm layer thickness and 30-50 µm pixel size). However, we posit that high-resolution DLP patterning can be achieved via further optimization of biocompatible resins and printing parameters.

Here, we develop an optimized photoresin based on fish gelatin and photoclick crosslinking chemistry for bioprinting perfusable, embedded microvascular networks via high-resolution digital light processing (DLP). Specifically, we created a protein-based photoresin composed of fish gelatin modified to enable thiol-norbornene photoclick chemistry (**Figure 1A**). This resin exhibits a low viscosity over a broad range of concentrations and temperatures.^26–29^ It is superior to synthetic-based resins (such as PEG-DA) due to its biocompatibility, biodegradability, and bioactivity. The resin is patterned using a custom-built DLP equipped with a digital micromirror device (DMD) capable of illuminating 4 million voxels per layer with a x-y resolution of ∼7.6 µm at the image plane and a z-step of 10 µm **(Figure 1A)**. We determined that pulsed (rather than continuous) illumination significantly improves printing resolution despite the negligible dark-curing associated with the thiol-norbornene photoclick chemistry. We note that printing in a pulsed mode likely reduces off-target crosslinking arising from the diffusion of excess initiating radical species. By introducing a biocompatible radical scavenger (DMPO), resin stability is improved yielding a longer printing window (>5 h), while the use of a biocompatible surfactant (Poloxamer-188) drastically reduced delamination and, hence, printing failures. Finally, as a simple demonstration, we printed hydrogels with embedded microvascular channels, including those with convoluted, multichannel, and multiscale geometries with minimum microchannel diameters of ∼20 µm.

**Figure 1.**
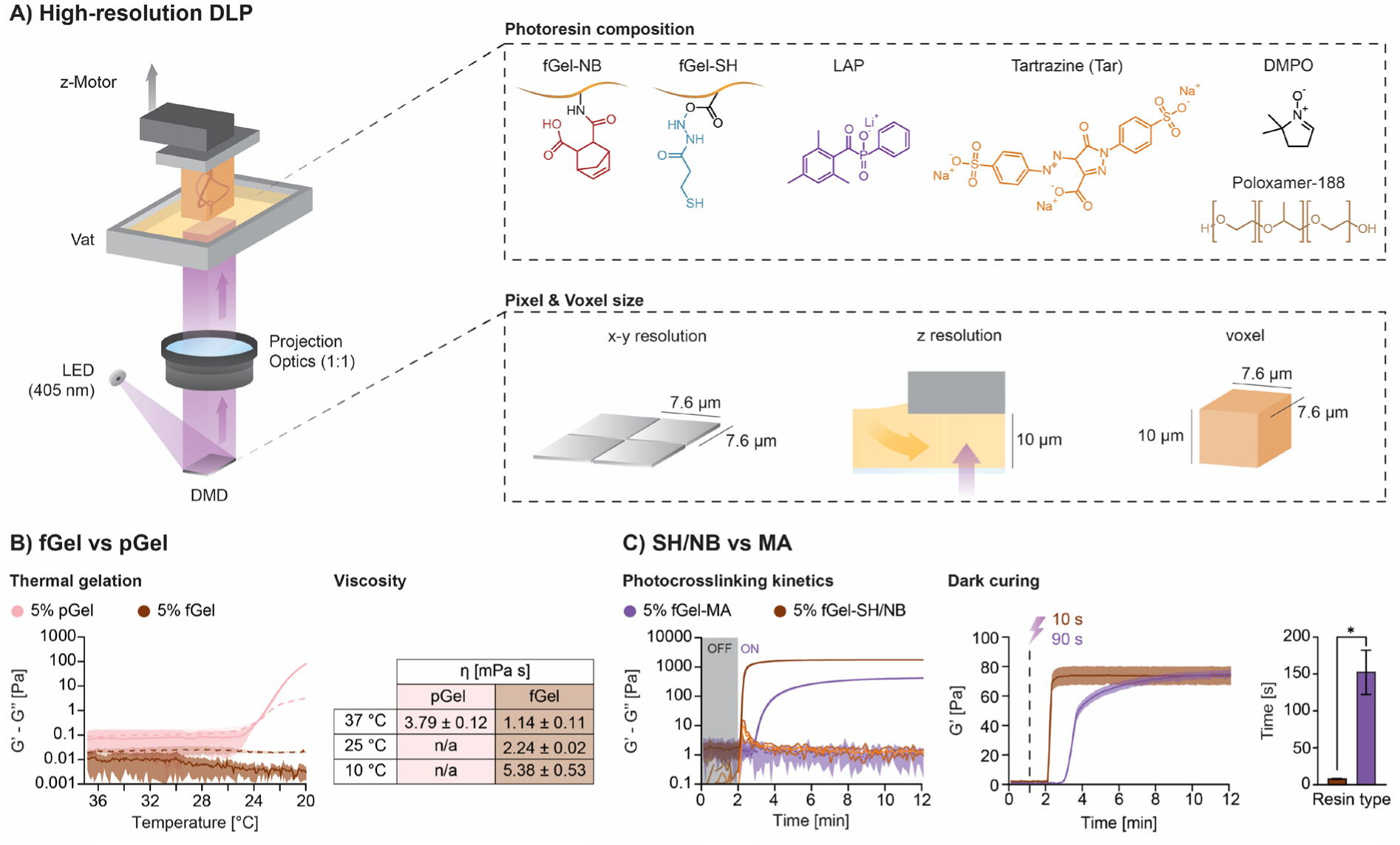
A) Schematic illustrations of DLP method and photopolymerizable resin moieties. B) Semi-log plot of shear moduli dependence on temperature (left) and apparent resin viscosity for porcine (pGel) and fish (fGel) gelatin (right). C) Semi-log plot of shear moduli dependence on time under UV (left) and dark (right) curing of methacrylated (fGel-MA) and thiol-norbornene fish gelatin (fGel-SH/NB) resins.

## 2. Results and Discussion

### 2.1. Photoresin design and optimization

Gelatin was chosen as the material platform over synthetic polymers for its known biocompatibility, biodegradability and presence of integrin binding sites that promote cell adhesion. As reported previously^26^, fish gelatin (fGel) offers improved thermal stability compared to conventional porcine gelatin (pGel).^27–29^ Due to its low proline and hydroxyproline content, fGel does not undergo thermal crosslinking upon cooling (**Figure 1B**). fGel solutions possess a remarkably low viscosity (∼0.002-0.003 Pa•s, **Figure 1B**), which is roughly 2-4 orders of magnitude lower than the viscosity range (∼0.25-10 Pa•s) of most commercial DLP resins. This low viscosity is necessary for resin flow in the narrow 10 µm gap between printed layers.

The fGel resin is modified with thiol (fGel-SH), norbornene (fGel-NB), and methacrylate (fGel-MA) pendant groups to enable photopolymerization (**Supplementary Figures S1-3**). Lithium phenyl-2,4,6-trimethylbenzoylphosphinate (LAP), the current gold standard for bioprinting, is used as a photoinitiator. We compared the photocrosslinking performance of thiol-norbornene-based and methacrylate-based resins with similar degree of substitution (∼0.19 mmolSH/NB or MA g-1), same polymer (5%) and LAP (0.05%) concentrations (**Figure 1C**). Photoclick fGel-SH/NB photoresin exhibited a significantly faster gelation onset and better crosslinking efficiency compared to fGelMA photoresin. Faster photocrosslinking kinetics are especially desirable when considering high-resolution applications that inherently require long printing times due to the thin slicing (10 µm) of 3D models. The step-growth nature of the thiol-norbornene chemistry also shows negligible dark curing compared to the traditional chain-growth mechanism of the methacrylate resin (**Figure 1C**). When the light is turned off, due to the chemical nature of the chain propagation mechanism, the fGel-MA photoresin continues its polymerization process in the dark, reaching a plateau G’ only after ∼2 min. This dark-curing phenomenon, as well as increased radical production and diffusion due to the required longer exposure times, is a source of off-target crosslinking and therefore a major drawback of this type of chemistry for high resolution applications. On the other hand, the step-growth nature of thiol-norbornene chemistry offers negligible dark curing (**Figure 1C**). Since the voxel resolution (7.6 x 7.6 x 10 µm^3^) benefits from the combination of efficient crosslinking, minimal excess radicals, and no dark curing, we chose fGel-SH/NB as the optimal material for our work. We note that thiol-norbornene hydrogels also exhibit better mechanical properties (lower shrinkage and brittleness) and biocompatibility compared to methacrylated hydrogels^30–32^. Given the large resin volumes needed to iteratively optimize the full set of printing parameters, we developed a one-step protocol for synthesizing fGel-NB^19^ (20 g batches) via a 1 h reaction, while fGel-SH (10g batches) is synthesized as previously reported.^31^

Next, we explored the effects of the degree of substitution (DS) and polymer concentration on the mechanical and bioactive properties of fGel-SH/NB hydrogels (**Figure 2**). First, we synthesized three different batches of fGel-SH/NB resins varying in their DS: ‘Low’ (∼0.09 mmol_SH/NB_ g^-1^), ‘Medium’ (∼0.14 mmol_SH/NB_ g^-1^), and ‘High’ (∼0.19 mmol_SH/NB_ g^-1^). Although the rapid nature of the click chemistry led to no significant difference in the gelation onset, the oscillatory photorheology measurements revealed that increasing DS leads to increased hydrogel stiffness (**Figure 2A**). Hydrogels with higher DS, and hence higher crosslinking density, exhibited lower swelling (**Figure 2A**). We also found that grafting the gelatin backbone with thiol and norbornene pendant groups did not alter its refractive index (**Supplementary Figure S4**). Notably, all fGel-SH/NB hydrogels synthesized are fully biodegradable. Each gel can be fully degraded by collagenase type IV (1 mg mL^-1^ solution) within 1 h of incubation (**Figure 2A**). This suggests that even at “High” DS, this endopeptidase can still recognize and hydrolyze the gelatin backbone, thus conferring the possibility for seeded cells to remodel the surrounding matrix. As shown by Wang et al.,^22^ resins susceptible to enzymatic degradation can be used to tune the mechanical properties of the final construct post-printing. We assessed the cell-adhesion properties of these hydrogels by seeding human umbilical vein endothelial cells (HUVECs) and observed that cell adhesion improves with higher DS (**Figure 2A, Supplementary Figure S5**).

**Figure 2.**
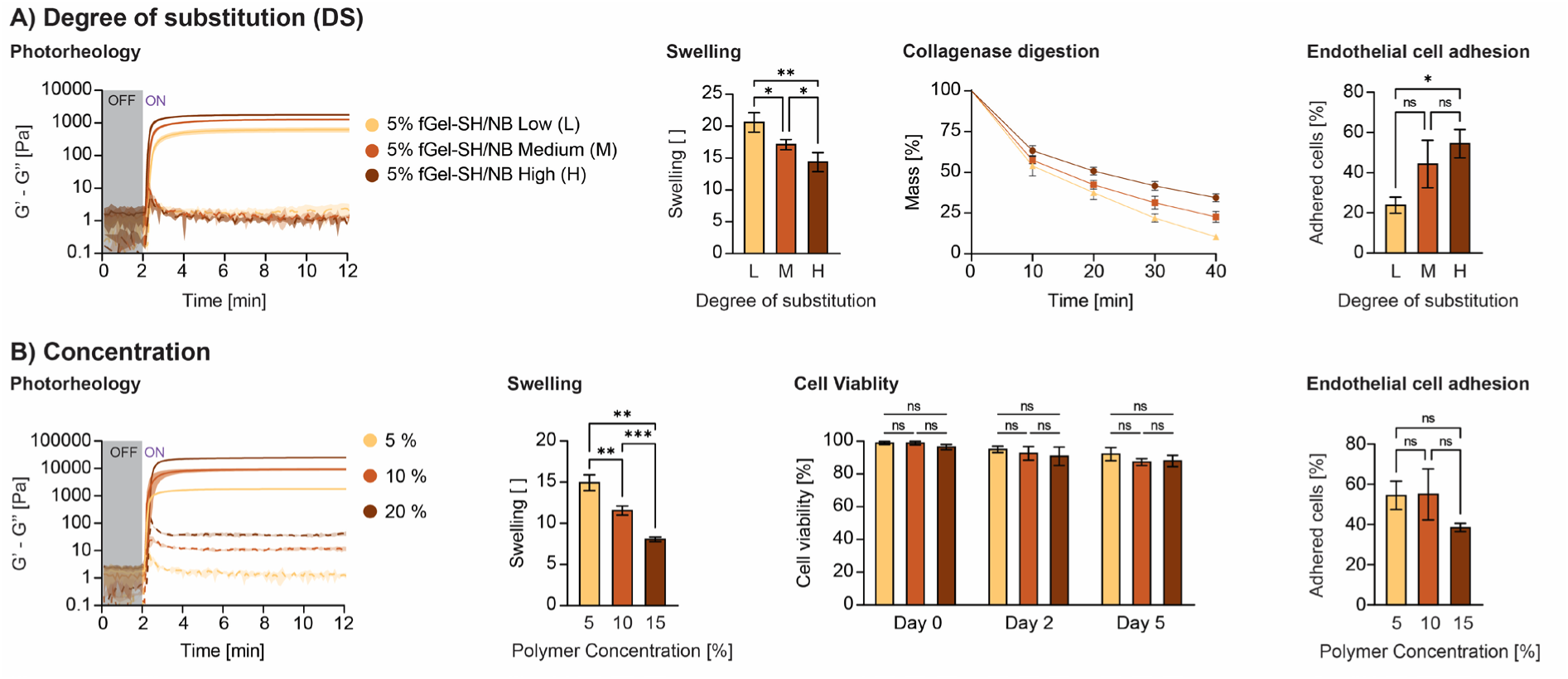
A) Photocrosslinking of fGel-SH/NB resins with various degrees of substitution (Low-L, Medium-M, and High-H). Swelling behavior, collagenase type IV digestion profile, and adhesion of human umbilical endothelial cells on hydrogel obtained with such resins. B) Photocrosslinking of fGel-SH/NB ‘High’ resins with various polymer concentrations (5%, 10%, and 20%). Swelling behavior, cell viability of embedded human neonatal dermal fibroblasts, and adhesion of human umbilical endothelial cells on hydrogel obtained with such resins.

We further modified the fGel-SH/NB with ‘High’ DS. Hydrogels with increasing polymer concentration, ranging from 5% to 20%, exhibited similar gelation onset, yet drastically increased shear elastic modulus (G’), ranging from 1772 ± 67 Pa (5% polymer) to 24954 ± 366 Pa (20% polymer), respectively (**Figure 2B**). Concomitantly, their degree of swelling decreased (**Figure 2B**), while their refractive index increased with increasing polymer concentration (**Supplementary Figure S4**). Next, human neonatal dermal fibroblasts (HNDFs) are incorporated to assess hydrogel biocompatibility. Excellent cell viability is observed after crosslinking (>95% at day 0) and during *in vitro* culture (>90% at day 2 and ≥ 87% at day 5) with no significant differences for different compositions (**Figure 2B**). Not surprisingly, confocal imaging revealed more pronounced cell spreading occurred in softer hydrogels (5-10% polymer) compared to the stiffest one (20% polymer) (**Supplementary Figure S6**). Interestingly, assessing the cell-adhesion properties by seeding HUVECs, we did not observe significant differences among the different conditions (**Figure 2B, Supplementary Figure S5**). Hence, we chose 10% fGel-SH/NB with ‘High’ DS as the resin of interest for high-resolution DLP printing screening.

### 2.2. Towards high-resolution printing

In DLP, the height resolution is determined by the printhead z-step and the ability of photoabsorber molecules to attenuate light penetration. Effectively blocking light from penetrating further into the resin is pivotal to avoid excessive “bleedthrough”, that is crosslinking over negative features such as fine perfusable microchannels running orthogonally to the light direction. First introduced by Benjamin et al.,^20^ tartrazine (Tar) has found widespread use in biocompatible photoresins for DLP printing.^8, 33–36^ This food dye has a molar extinction coefficient three orders of magnitude higher than LAP at the irradiation wavelength (405 nm) (**Supplementary Figure S7**), and thus acts as an effective photoabsorber. Tests under oscillation photorheology show that even at low concentration (0.025-0.1%) Tar drastically affects the gelation speed of the fGel-SH/NB photoresin in a dose-dependent manner (**Supplementary Figure S8A**). Interestingly, within the tested concentration range, Tar does not affect the refractive index of the resin (**Supplementary Figure S8A**). To counterbalance the Tar-induced gelation delay, one can increase the photoinitiator (LAP) concentration (**Supplementary Figure S8B**). At relatively high LAP concentrations (0.25-1%), there is a slight increase in the refractive index of the photoresin (**Supplementary Figure S8B**). Importantly, the Tar-induced gelation delay also depends on the selected photorheology gap. Since light intensity decreases exponentially as it penetrates through the resin (Lambert-Beer Law), the difference in gelation time observed for varying Tar concentration is far more pronounced when tests are carried out using larger gap thicknesses (i.e., 200 µm vs 50 µm **Supplementary Figure S9A**). Increasing Tar concentration from 0.05% to 0.2% is necessary to achieve a sufficient gelation time difference between these gap thicknesses and ensure minimal bleedthrough during printing process with a 50 µm layer height. Higher Tar concentration is likely necessary to achieve a 10 µm layer height. Besides increasing the LAP content (**Supplementary Figure S8B**), another strategy to compensate for the Tar-induced gelation delay is to increase the polymer concentration (**Supplementary Figure S9B**).

To further assess this resin formulation for DLP printing, we printed 4x4x4 mm cubes with an open microchannel composed of vertical, horizontal, and curved parts. The printed cubes are washed in PBS and injected with a red dye filler for perfusion assessment and imaging (**Supplementary Figure S10**). Using a resin composed of 10% fGel-SH/NB, 0.15% Tar, and 0.5% LAP, we first screened the effects of varying light intensity (10, 20, 30, and 40 mW cm^-2^) and irradiation mode (continuous vs. pulsed). We observed that 20 mW cm^-2^ in a pulsed setting is sufficient to embed microchannels that are partially perfusable with these hydrogels. Next, we increased the Tar concentration (up to 0.375%) to minimize bleedthrough. High Tar concentrations resulted in print failure due to delamination (low cohesion between the layers), which was ameliorated by increasing LAP (0.6%) and polymer (20%) concentrations. Finally, we increased the layer exposure time from 8 s to 12 s to achieve stable, perfusable microchannels that could withstand dye injection without deformation or defect formation (e.g., bulging or infiltration between poorly adhered layers) (**Supplementary Figure S10**).

Importantly, pulsed illumination appears to be critical for printing hydrogels with fine microchannels (**Figure 3A**). Continuous illumination at 10 mW cm^-2^ led to delamination due to poor layer adhesion arising from insufficient light penetration and free radical generation. Upon increasing the light intensity to 20 mW cm^-2^, the penetration depth and free radical generation increases resulting in clogged microchannels. After screening different pulsed illumination sequences (**Supplementary Figure S11**), we find that on/off pulses of 16.5 ms in duration each result in the best print fidelity. Longer “on” states resulted in visible clogging, while shorter ones resulted in striation defects associated with poor crosslinking. By dividing the energy deposited in the resin in discrete and interspersed packages, we limit the formation and diffusion of radical excess allowing the crosslinking reaction to consume free radicals generated during the “off” mode prior to forming new free radicals during the “on” pulse. To further reduce build failures, we lowered the resin surface tension from 67.0 ± 0.7 mN m^-1^ to 59.5 ± 2.4 mN m^-1^ by adding 0.1% Poloxamer-188 (Pol-188), a biocompatible non-ionic surfactant (**Supplementary Figure S12**). This resin modification reduced the observed printing failures due to delamination from 30% (without surfactant) to nearly 0% (with surfactant).

**Figure 3.**
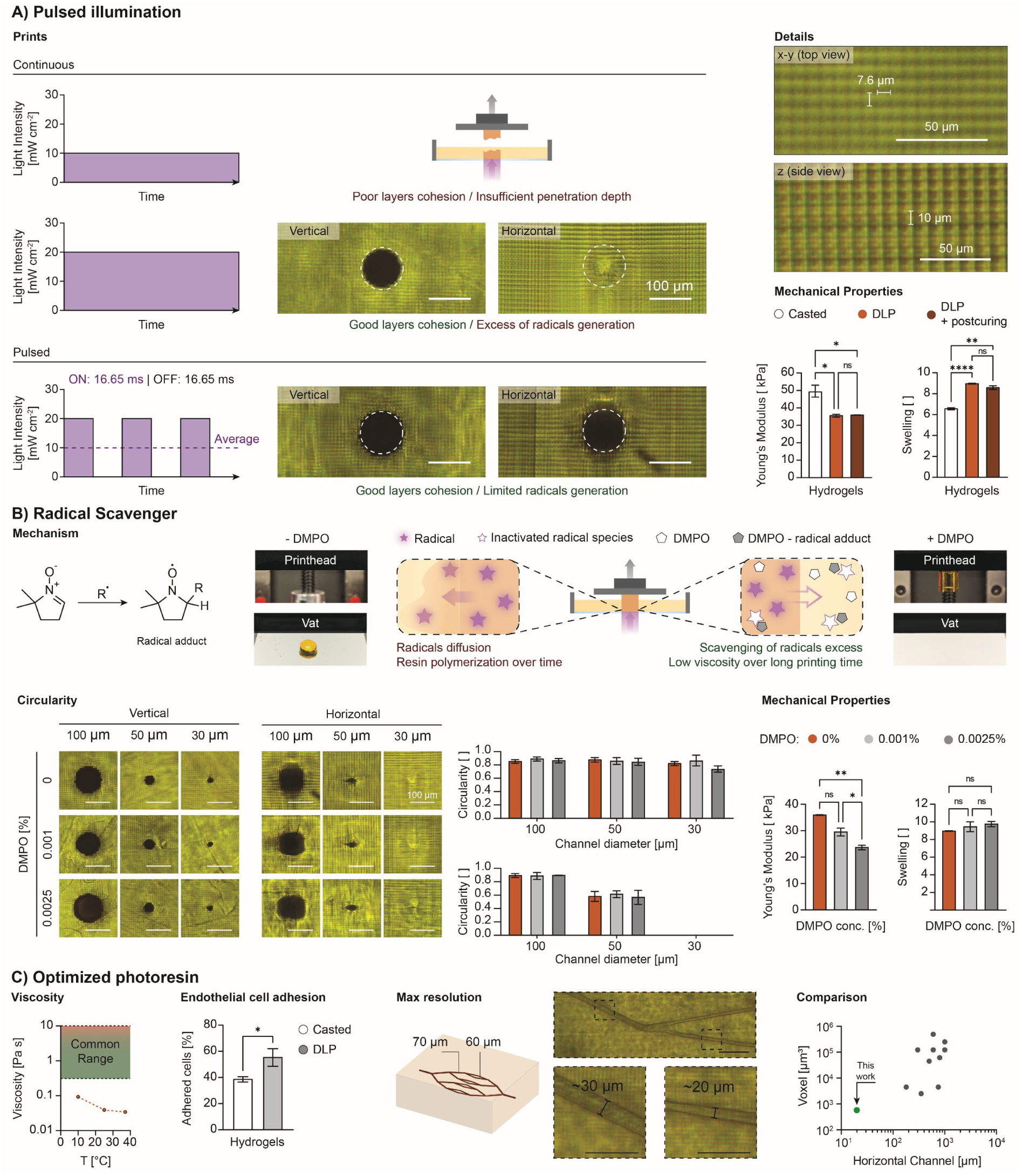
A) Continuous vs pulsed illumination patterns (left), details of x-y pixels and layers on printed parts, and mechanical properties of hydrogels obtained via casting vs DLP printing vs DLP printing + postcuring (right). B) Scavenging mechanism of DMPO and examples of long prints (5 h 45 min, 1 cm height) with and without it (top). Circularity of microchannel openings (left) and mechanical properties of hydrogels printed with and without DMPO (right). C) Viscosity of the optimized photoresin (left), and adhesion of human umbilical endothelial cells on hydrogel obtained with such resin (middle). Details of smallest perfusable microchannels and comparison with recent work on DLP printing with biocompatible hydrogel precursors (right). [dots will be changed to reference number when we have a final version of the manuscript] Scale bars: 100 µm.

Hydrogels printed using a resin formulation composed of 20 % fGel-SH/NB, 0.6% LAP, 0.375% Tar, and 0.1% Pol-188 had a lower Young’s modulus (35.9 kPa) under uniaxial compression compared to the cast hydrogels without Tar (49.6 kPa) (**Figure 3A**) and slightly higher degree of swelling. Interestingly, possibly due to the high Tar concentration which significantly reduces light penetration, we did not observe any differences in compression modulus and swelling degree after post-curing the printed hydrogels for 20 min under 405 nm illumination (20 mW cm^-2^) (**Figure 3A**). New failure modes are observed when printing larger hydrogel constructs. In particular, the performance enhancement associated with surfactant additions is drastically reduced for longer print times (5 h in duration, >8.5 mm thick constructs). We hypothesize the slow polymerization of the resin in the vat due to excess radical diffusion may lead to the hydrogel adhering more to the vat than the build plate (**Figure 3B, Supplementary Figure S13**). A potential solution is represented by radical scavengers, such as 2,2,6,6-tetramethyl-1-piperidinyloxy (TEMPO), which has been recently used to introduce non-linear gelation threshold response in a thiol-ene resin (allyl ethers as -enes).^37^ Interestingly, when TEMPO is used within the previously reported concentrations (0.001-0.01%), we did not observe any significant difference in gelation onset in our resin (**Supplementary Figure S14A**). Hence, we investigated 5,5-Dimethyl-1-pyrroline N-oxide (DMPO) as an effective radical scavenger for thiol-norbornene resins in light-mediated 3D printing (**Figure 3B**). DMPO, which has higher water solubility and biocompatibility,^38^ significantly delays the onset of gelation (**Supplementary Figure S14B**). Adding DMPO to the resin formulation did not compromise the print resolution, i.e., microchannels as small as 30 µm along the z axis (light direction) and 50 µm in the x-y plane (orthogonal to the light direction) (**Figure 3B**) could be fabricated. Vertically printed microchannels are characterized by a high circularity (∼0.7-0.9) across several diameters tested (100, 50 and 30 µm). On the other hand, printing negative features on the x-y plane unavoidably leads to a reduction in resolution due to the bleedthrough required for layers cohesion. The horizontally printed microchannels showed a high circularity (∼0.85-0.9) for 100 µm microchannels, a drastic reduction in circularity (∼0.5) for the 50 µm microchannels and occluded 30 µm microchannels. While these results suggest that it would be possible to create small cylindrical horizontal microchannels using an even higher resolution DLP (smaller z-step), the ability to seed endothelial cells of diameter ∼10-20 µm within such capillary-like microchannels is quite difficult. It should be noted that DMPO interferes with the free-radical crosslinking process leading to a modest reduction in hydrogel stiffness (**Figure 3B**) and increased swelling behavior (**Figure 3B**). However, its incorporation vastly decreased printing failures when patterning thick hydrogels (∼ 1 cm) with embedded microchannel networks (**Supplementary Figure S13**).

The final optimized resin formulation (20 % fGel-SH/NB, 0.6% LAP, 0.375% Tar, 0.1% Pol-188, 0.001% DMPO) exhibits a low viscosity across a broad range of temperatures (i.e., 34.26 ± 0.55, 39.03 ± 2.18, and 92.85 ± 6.00 mPa s at 37°C, 25°C and 10°C, respectively (**Figure 3C**). These measured viscosity values are roughly 10-100x lower than commercial DLP resins. Its refractive index (1.3654) is within the range of values reported for cells and cytoplasmic organelles (1.36-1.39)^9^ (**Supplementary Figure S15**). The adhesion of endothelial cells (HUVECs) on DLP printed constructs is higher than the cast controls (55.2 ± 5.5 % vs 38.5 ± 1.6 %) (**Figure 3C**). No significant changes in microchannel diameter are observed upon 24 h swelling in PBS (**Supplementary Figure S16).**

Upon perfusion of printed microchannels, we observed a mismatch between the target (100 µm) and the experimental diameter (**Supplementary Figure S17**). While this mismatch falls between a ± 2 pixels error margin for the vertical prints (∼6-10 µm), it significantly increases for the horizontal prints (∼32-38 µm). The bleedthrough necessary to achieve layer cohesion, the limited flow of resin in small microchannels and the larger z-step all contribute to this mismatch. Importantly, accounting for this discrepancy, one can print the desired microchannel size in the horizontal plane. Indeed, perfusable microchannels of ∼20 µm in diameter have been achieved (**Figure 3C**). These dimensions, once accessible in biocompatible hydrogels solely by 2P-SL,^10, 11, 17^ are now achievable via DLP (**Supplementary Table S1**).

### 2.3. Embedding Complex Microchannel Networks

Using high-resolution DLP, we printed hydrogels with embedded microchannel networks composed of one or two spiral microchannels (500 µm diameter), which are perfused with red- and blue-dyed fillers (**Figure 4A**, left, middle). We also printed a hydrogel in which two spiral microchannels are wrapped around a central horizontal microchannel (evacuated / air filled) to further highlight these capabilities (**Figure 4A**, right). We also embedded a glomerular-like capillary network within a printed hydrogel construct. (**Supplementary Figure S18**).

**Figure 4.**
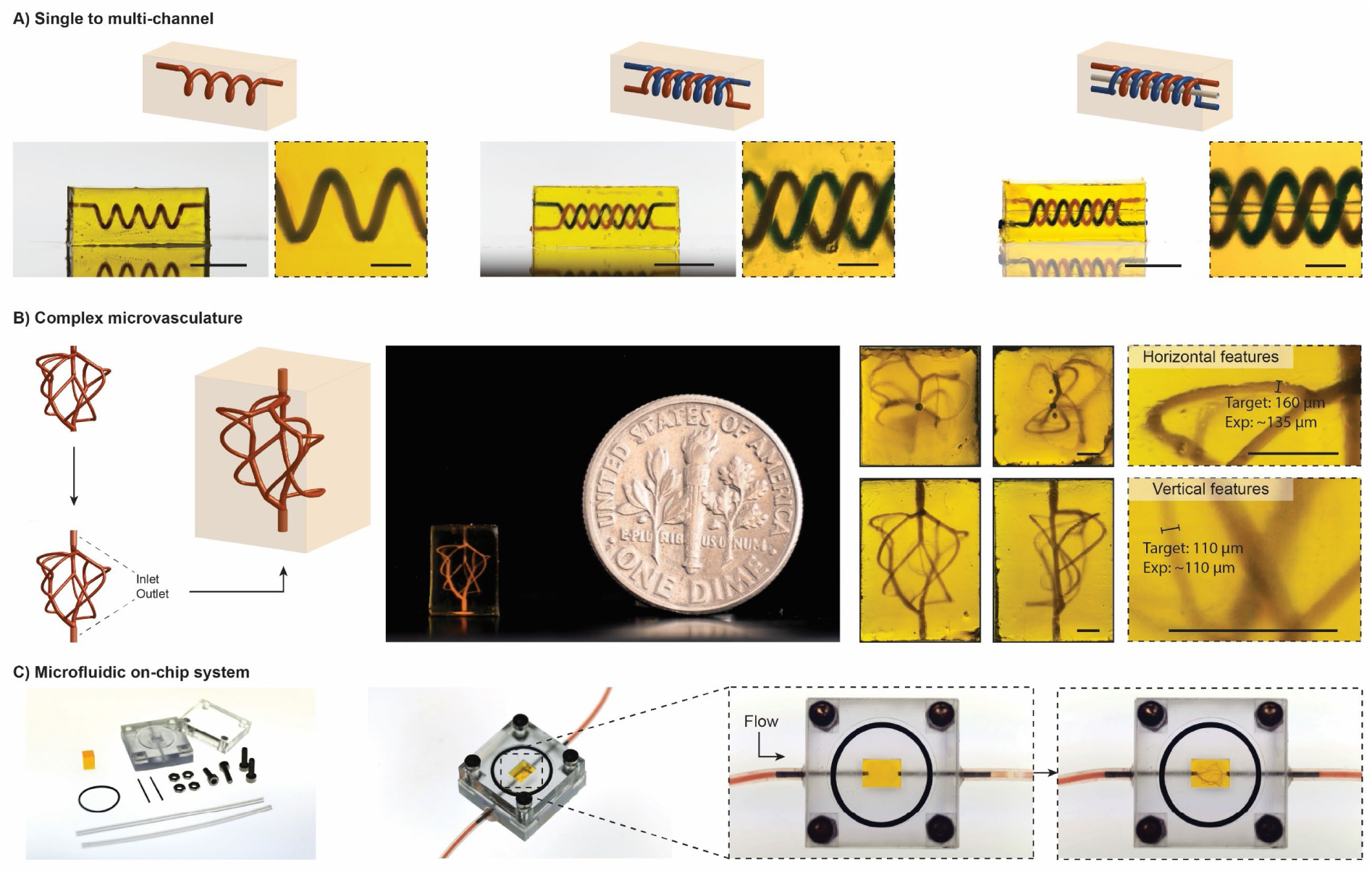
A) Printed hydrogels with embedded single (left) or multiple (middle-left) perfusable microchannels. Scale bars: 4 mm. B) Schematic of vascular model design (left). Picture (middle) and microscopy close-ups (right) of a 6 mm tall construct featuring biomimetic multiscale microvasculature. Scale bars: 100 µm. C) Compatibility with microfluidic on-chip system.

As a final demonstration, we used a recently reported algorithm to generate a biomimetic vascular tree.^39^ This microvascular network, which features 15 branching points (**Figure 4B**), we embedded within hydrogel constructs printed with thicknesses of 6 mm, 4.8 mm, and 3.6 mm, respectively, and perfused using dyed filler (**Figure 4B, Supplementary Figure S19**). There is good agreement between the target and printed microchannel sizes as small as 70 µm in diameter. To further demonstrate their utility, we placed these printed hydrogel constructs with embedded microvascular networks in a custom-designed chip and demonstrated their perfusion with dyed filler (**Figure 4C**).

## 3. Conclusion

In summary, we reported an optimized biocompatible photoresin and pulsed light irradiation sequence that enables high resolution DLP printing of hydrogel constructs with embedded microchannels as well as microvascular networks. We have characterized the effects of compositional modifications on their resin rheology, printing resolution, and hydrogel stiffness and swelling behavior. We have also shown that such constructs present high cell biocompatibility and good adhesion properties towards human endothelial cells. Our platform is well suited for printing complex organ-on-chip models for myriad biotechnology applications.

## 4. Experimental Section

### Thiolated fish gelatin synthesis (fGel-SH)

fGel-SH (High DS) was synthesized based on the protocol from Rizzo *et al.*^31^ Briefly, 10 g of gelatin from cold water fish skin (Sigma-Aldrich) were dissolved in 500 mL of 150 mM MES buffer pH 4.5. Next, 0.3 g of 3,3’-dithiobis(propionohydrazide) (DTPHY, Toronto Research Chemicals) was added to the solution. Once completely dissolved, 1.5 g of 1-ethyl-3-(3-dimethylaminopropyl)carbodiimide (EDC, TCI America) was added and the reaction was left to proceed for 24 h. 1.7 g of TCEP was then added, and the reduction of disulfides was left to proceed for 8 h under gentle stirring. After addition of 1 g of sodium chloride (NaCl), the solution was dialyzed (MWCO: 14 kDa, Ward’s Science) against acidified mQ H_2_O (pH 4 using diluted HCl) for 4 days with frequent water changes and freeze-dried. Lyophilized fGel-SH was stored under inert atmosphere at −20 °C prior to use. The degree of substitution (DS) was estimated to be ∼0.18 mmol of SH per gram of fGel with ^1^H-NMR in D_2_O using internal standard 3-(trimethylsilyl)-1-propanesulfonic acid (DSS, 2H, ∼0.65 ppm) and hydrazide methylene protons (∼2.7 and 2.8 ppm, see **Supporting Figure S1**).

### Fish gelatin norbornene synthesis (fGel-NB)

fGel-NB (High DS) was adapted from previously reported protocols.^40, 41^ Briefly, 20 g of gelatin from cold water fish skin (Sigma-Aldrich) were dissolved in 200 mL of 1 M carbonate-bicarbonate buffer pH 9. When completely dissolved, 2.5 g of cis-5-norbornene-endo-2,3-dicarboxylic anhydride (CA) was added to the reaction mixture under vigorous stirring and the reaction was left to proceed for 1 h. Next, the solution was diluted 2-fold with milliQ H_2_O, filtered using a 0.22 µm filter, dialyzed (MWCO: 14 kDa, Ward’s Science) against mQ H_2_O for 4 days with frequent water changes and finally freeze-dried. The degree of substitution (DS) was estimated to be ∼0.19 mmol of NB per gram of fGel with ^1^H-NMR in D_2_O using internal standard 3-(trimethylsilyl)-1-propanesulfonic acid (DSS, 2H, ∼0.65 ppm) and NB protons (∼6.4–6.0 ppm, see **Supporting Figure S2**).

### Methacrylated fish gelatin synthesis (fGel-MA)

1 g of gelatin from cold water fish skin (Sigma-Aldrich) was dissolved in 10 mL of 0.1 M carbonate-bicarbonate buffer pH 9. When completely dissolved, 0.3 mL of methacrylic anhydride was added to the reaction mixture under vigorous stirring and the reaction was left to proceed for 1.5 h. Next, the mixture was centrifuged at 4000 rpm for 5 minutes. 0.5 g of sodium chloride (NaCl) was added to the supernatant, and the solution was dialyzed (MWCO: 14 kDa, Ward’s Science) against mQ H_2_O for 4 days with frequent water changes and freeze-dried. Lyophilized fGel-SH was stored under inert atmosphere at −20 °C prior to use. The degree of substitution (DS) was estimated to be ∼0.20 mmol of MA per gram of fGel with ^1^H-NMR in D_2_O using internal standard 3-(trimethylsilyl)-1-propanesulfonic acid (DSS, 2H, ∼0.65 ppm) and MA protons (∼5.45 and 5.70 ppm, see **Supporting Figure S3**).

### Absorption spectra

0.7 mL of each solution was pipetted in quartz glass cuvettes (1 cm optical path, Thorlabs) and analyzed at 25°C, with 1 nm step, using a BioTek Synergy HT reader equipped with Take3 microplate, and subtracting PBS (or H_2_O for IDX) background.

### Photorheology

Photorheology was carried out on a Discovery HR 20 (TA instruments) rheometer equipped with an 8 mm parallel plate stainless steel geometry and quartz glass floor. Omnicure Series2000 lamp (Lumen Dynamics) with 400-500 nm filter was used as light source. Light intensity at 405 nm was measured using a photodiode power sensor (S140C, Thorlabs) and a 410 nm (FWHM: 10 nm) bandpass filter (Thorlabs) placed at the sample location. The photoresins were prepared by mixing and diluting in PBS pH 7.4 the stock solutions of the various components to obtain the desired final concentrations. Oscillatory measurements were performed in triplicate using 13 µL of photoresin with a 200 µm gap (unless otherwise stated), at 1 Hz frequency, and 1.5% strain. All tests were conducted in the dark, and in the presence of a water-soaked tissue to prevent samples from drying. Light irradiation was initiated after 120 s.

### Viscosity measurements

Viscosity was measured on a Discovery HR 20 (TA instruments) equipped with a double-gap geometry. Flow sweep measurements were performed with a shear rate ranging from 0.1 to 100 s^-1^ using 12 mL of the tested solutions. After reaching the desired temperature, samples were left to equilibrate for 5 min before starting the measurement. Reported values refer to 1 s^-1^.

### Swelling tests

Photoresins were prepared as described above, cast in PDMS rings (6 mm inner diameter, 2 mm height) and crosslinked for 5 min with 405 nm irradiation (10 mW cm^-2^). Crosslinked hydrogels were left to swell to equilibrium for 24 h in PBS (6 mL per gel) at room temperature under gentle shaking. Next, the samples (n = 5) were weighted (hydrated mass) and then freeze-dried to obtain the dry mass. Swelling ratio was then calculated as ratio between hydrated and dry mass.

### Compression tests

Photoresins without Tar were cast in PDMS rings (4 mm inner diameter, 2 mm height) and crosslinked for 5 min with 405 nm irradiation (10 mW cm^-2^). Crosslinked hydrogels were left to swell to equilibrium for 24 h in PBS (6 mL per gel) under gentle shaking. Cylinders of the same dimensions were DLP printed using photoresins with 0.35% Tar, and left to swell to equilibrium in PBS for 24 h. The DLP “post-cured” samples were instead exposed to 405 nm irradiation (20 mW cm^-2^) in a UV box for 20 min immediately after printing, and then left to swell in PBS. Images of the swollen gels were taken with a stereomicroscope (Discovery V.20, Zeiss) and used to measure the surface area on Fiji ImageJ. Compression tests were carried out on a Discovery HR 20 (TA instruments) equipped with an 8 mm stainless steel flat geometry. Swollen samples were placed between the plates and compression was performed at 10 µm s^-1^. Region between 0-5% strain was used to calculate the Young’s modulus.

### Refractive index (RI) measurements

RI was measured using a digital refractometer (81150-56, Cole-Parmer). Prior to the measurements, the instrument was zeroed using milliQ H_2_O.

### Pendant drop

Surface tension was measured at room temperature via the pendant drop method following protocol by Daerr et al. using a stainless-steel blunt nozzle (1.83 mm OD, 14 Gauge, Nordson).^42^ Images of stable hanging drops were analyzed with ImageJ using the open-source Pendant_Drop plugin.^42^

### Collagenase digestion

Photoresins composed of 5% fGel-SH/NB (Low, Medium, and High DS) and 0.05% LAP in PBS were cast in PDMS rings (4 mm inner diameter, 1.5 mm height) and crosslinked in a UV box for 5 minutes (10 mW cm^-2^). The crosslinked hydrogels were left to equilibrate for 24 h in PBS (6-well plate under gentle orbital shaking). Hydrogels were weighed (time 0) and then placed in 0.5 mL solution of 1 mg mL^-1^ Collagenase Type IV (StemCell Technologies) in an incubator at 37 °C. Samples (n = 6) were weighed after 10, 20, 30 and 40 min to monitor the weight loss via enzymatic degradation. All hydrogels were found to be fully degraded after about 60 min.

### Cell culture

Foreskin human fibroblasts (NHDF, passage 7-9) were cultured in DMEM with 10% v/v FBS and antibiotic-antimycotic (100 units mL^-1^ penicillin, 100 µg mL^-1^ streptomycin, and 0.25 µg mL^-1^ Gibco Amphotericin B) with media change every other day. HUVECs (passage 3-7) were cultured in EGM-2 (Lonza) with media change every other day. For passaging cells were detached using 0.05% (for HUVECs) and 0.25% (for NHDF) trypsin.

### Live/Dead assay and phalloidin staining of cast gels

NHDF were resuspended at 0.5 million mL^-1^ in photoresins composed of 5%,10, and 20% fGel-SH/NB (High DS) and 0.05% LAP, cast in PDMS rings (4 mm inner diameter, 1.5 mm height), and crosslinked in a UV box for 5 minutes (10 mW cm^-2^). Samples were cultured for 5 days with media change every other day. Cells-embedded hydrogels (n = 3 per time point) were taken on day 0, day 2, and day 5 and incubated in FluroBrite^TM^ DMEM supplemented with 1:2000 calceinAM and 1:500 ethidium homodimer-1 for 30 mins. Imaging (200 µm, 10 µm steps) was performed on a confocal microscope (Imager Z2, Zeiss) equipped with a water-immersion objective (10x/0.3). Cell viability was quantified by counting viable (calcein) and dead (ethidium homodimer-1) cells with the ImageJ Analyze particle function. For phalloidin staining, samples on day 5 were fixed in 4% paraformaldehyde for 30 min at room temperature and stained using ActinGreen^TM^ 488 ReadyProbes^TM^ and DAPI (30 min incubation followed by 3x washing in PBS). Images were acquired on a confocal microscope (LSM 980, Zeiss).

### HUVECs adhesion tests

Cylindrical cast and DLP printed gels were prepared as described above. After 24 h swelling in EGM-2, the media was removed and 20 µL HUVEC suspension (0.3 million cells mL^-1^ in EGM-2) was pipetted onto the samples surface. Cells were left to sediment and adhere for 1.5 h. After 3x washes with PBS, the samples (n = 3) were incubated in FluroBrite^TM^ DMEM supplemented with Hoechst 33342 (NucBlue^TM^ Live ReadyProbes^TM^) for 30 mins. Imaging of the nuclei of adhered cells was performed on a confocal microscope (Imager Z2, Zeiss) equipped with a water-immersion objective (10x/0.3). Images were analyzed with ImageJ and percentage of adhered cells was estimated based on the total number of cells pipetted onto the gel surface.

### DLP (bio)printing

Printing was performed on a MONO3-VZ1 (Monoprinter, part of Microfluidics for all, Inc.) equipped with an LRS-WQ SL LC UV projection core (Visitech) featuring a digital micromirror device with 2560x1600 pixels (7.56 µm pixel size). 1.5 - 4 mL of photoresin formulations, prepared as previously described, were sterile filtered (0.22 µm Millex^®^-GP, Millipore Sigma) and transferred to the DLP vat. Prints were performed with 10 µm z-step, 1 mm lifting height, 0.5 mm s^-1^ printhead speed, and 2 s pre-exposure holding time.

### 3D models

The biomimetic microvasculature model **(Figure 4B, Supplementary Figure S19)** was generated using a public python package (https://github.com/zasexton/Synthetic-Vascular-Toolkit) based on the work of Sexton et al.^39^ Inlet and outlet were added to the resulting vascular network and the model was then subtracted from a cuboid to result in an embedded perfusable vascular tree. For chip-perfusion test (**Figure 4C**), the construct was left to swell for 24 h, and then placed into the custom designed chip printed via sterolithography (Form3B, FormLabs) using BioMed Clear resin. Watertight seal was obtained using O-rings (McMaster-Carr), which were compressed via screwing a laser-cut acrylic plates (McMaster-Carr). Inlet and outlet pins were glued (Epoxy, Loctite) to the main body of the chip. The glomerulus construct (**Supplementary Figure S18**) was similarly designed by adapting an existing model (Turbosquid 2132728, Shutterstock Inc.). All other 3D models were designed using Fusion360, Solidworks or Onshape. Perfusability was tested with injection of Micorfil® (Flow Tek, Inc.).

All chemicals were purchased from Merck, and cell culture reagents from ThermoFisher Scientific unless otherwise specified.

### Statistical analysis

Statistical analysis was performed using GraphPad Prism 9 and statistical significance was determined using one-way Welch Anova with multiple comparisons, two-way Anova with multiple comparisons or unpaired Welch’s t-test.

## Competing Interest Statement

J.A.L. serves on the Scientific Advisory Board of Azul 3D and Trestle Biotherapeutics.

## Acknowledgements

The authors gratefully acknowledge support from the Vannevar Bush Faculty Fellowship Program sponsored by the Basic Research Office of the Assistant Secretary of Defense for Research and Engineering through the Office of Naval Research Grant N00014-21-1-2958 and the National Science Foundation through CELL-MET ERC (#EEC-1647837). R.R. acknowledges Swiss National Science Foundation Postdoc.Mobility fellowship (P500PM_214191). We thank the Harvard John A. Paulson School of Engineering and Applied Sciences Molecular and Cellular Biology Core (SEAS MCB) for infrastructure and support. We thank the Harvard Center for Biological Imaging (RRID:SCR_018673) for infrastructure and support.

