## Supporting Information for "Embedding Perfusable Microchannel Networks in Photoclickable Bioresins via High-Resolution Digital Light Processing"

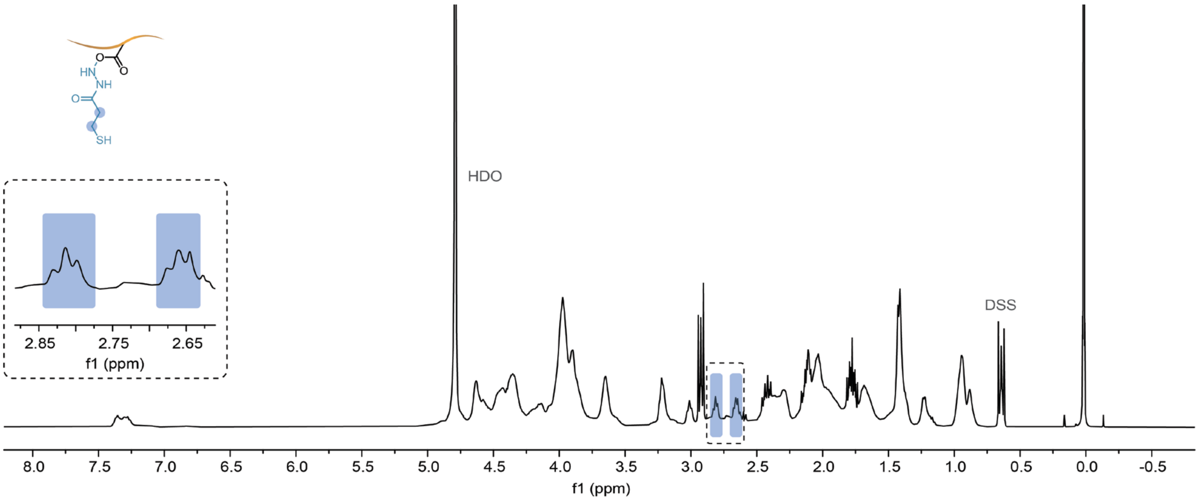


**Figure S1.** Thiolated fish-gelatin (fGel-SH) $^1$H-NMR.

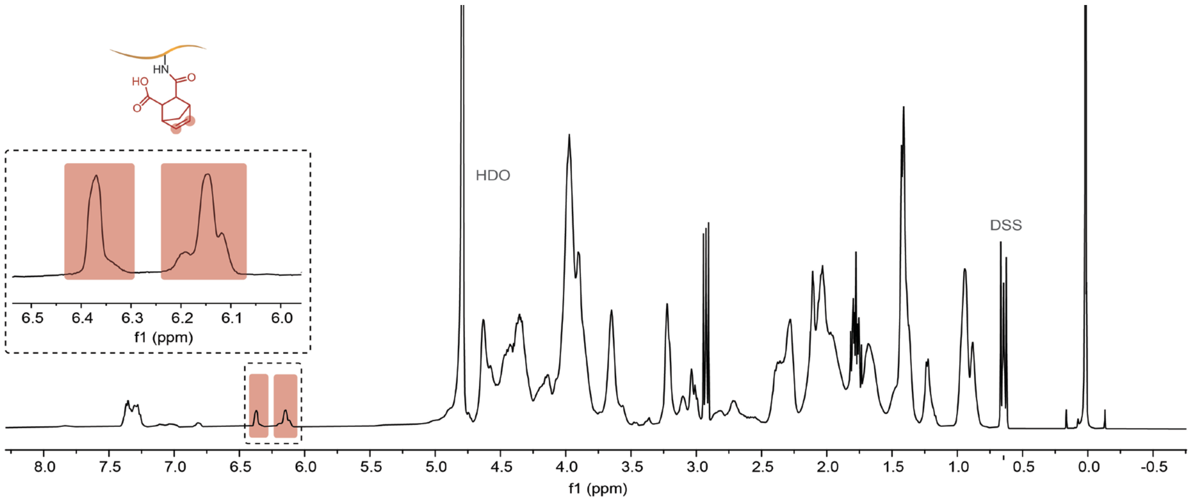


**Figure S2.** Fish-gelatin norbornene (fGel-NB) $^1$H-NMR.

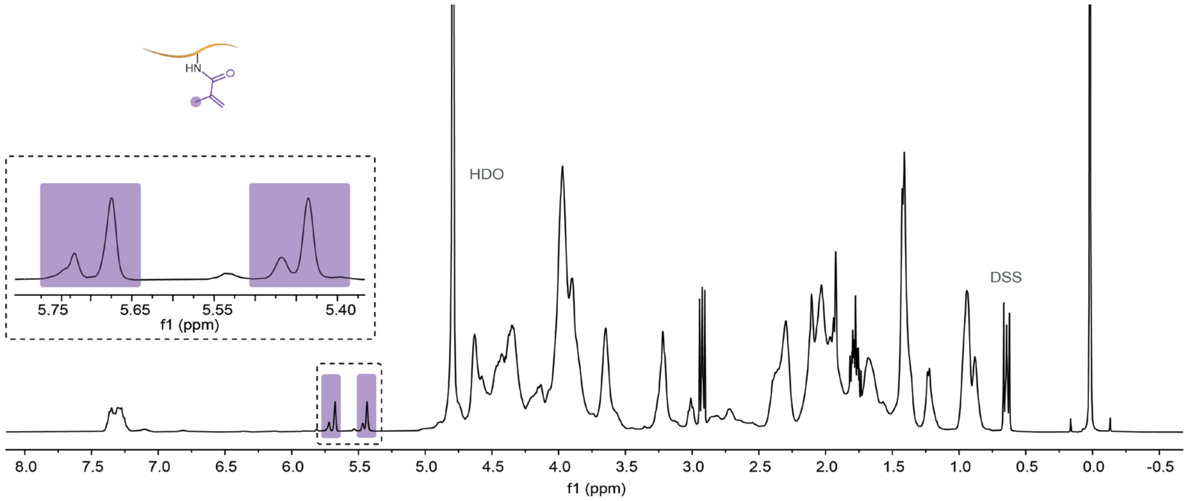


**Figure S3.** Fish-gelatin methacrylate (fGel-MA) $^{1}$H-NMR.

**A) Degree of substitution (DS)**

| | RI |
|---|---|
| Low | 1.3430 |
| Medium | 1.3430 |
| High | 1.3430 |

**B) Concentration**

| | RI |
|---|---|
| 5% | 1.3430 |
| 10% | 1.3506 |
| 20% | 1.3654 |

**Figure S4.** A) Refractive index (RI) of 5% fGel-SH/NB with various degree of substitution (Low: ~ 0.09 mmol$_{SH/NB}$ g$^{-1}$, Medium: ~ 0.14 mmol$_{SH/NB}$ g$^{-1}$, and High: ~ 0.19 mmol$_{SH/NB}$ g$^{-1}$). B) Refractive index (RI) of fGel-SH/NB 'High' at various polymer concentrations.

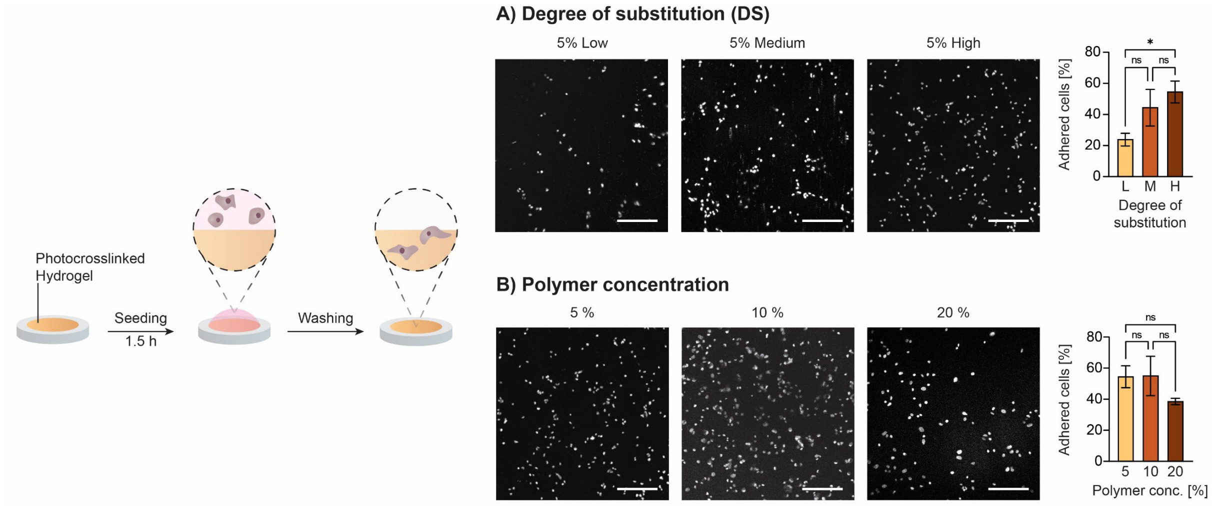


**Figure S5.** Endothelial cell (HUVECs) adhesion on cast hydrogel constructs produced using fGel-SH/NB photoresins with (A) varying degrees of substitution (Low-L, Medium-M, High-H) and (B) polymer concentrations. Confocal images show nuclei staining (grey). Scale bars: 200 μm.

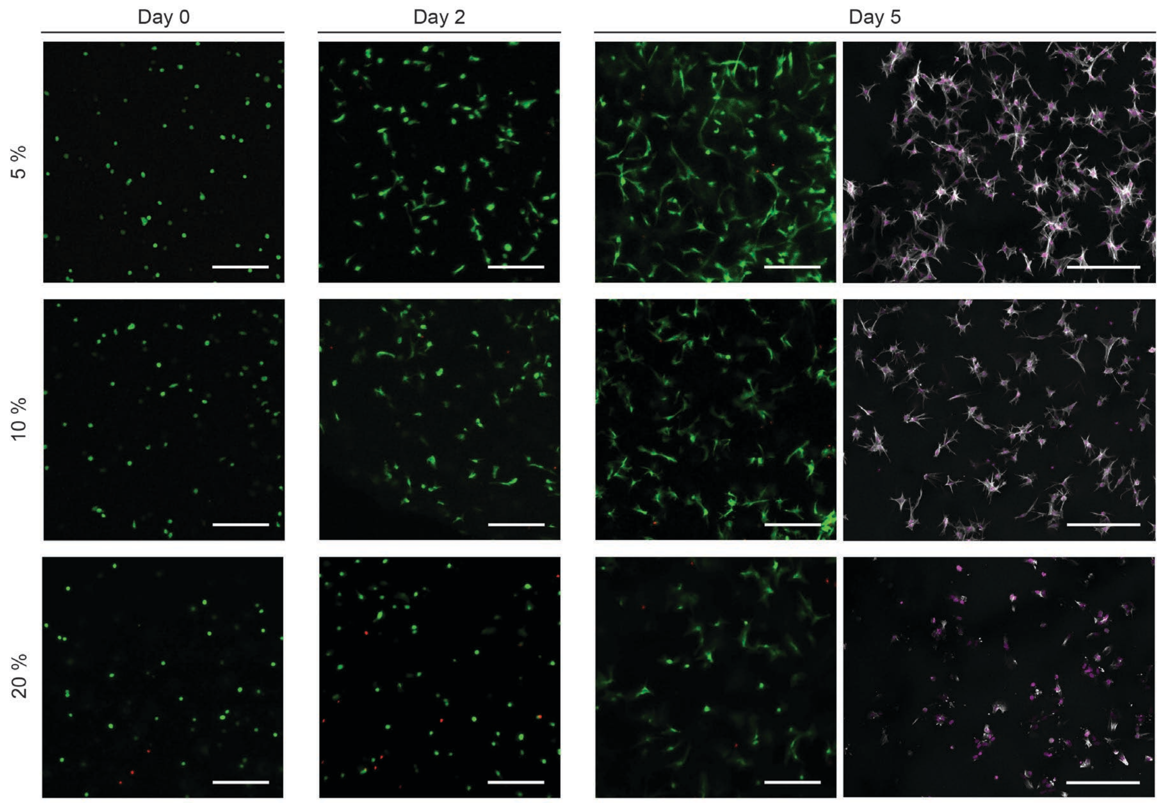


**Figure S6.** Live/dead assay of human neonatal dermal fibroblasts embedded in fGel-SH/NB 'High' photoresin at various polymer concentrations. On the right column, actin (grey) and DAPI (magenta) staining show cells morphology at day 5. Scale bars: 200 µm.

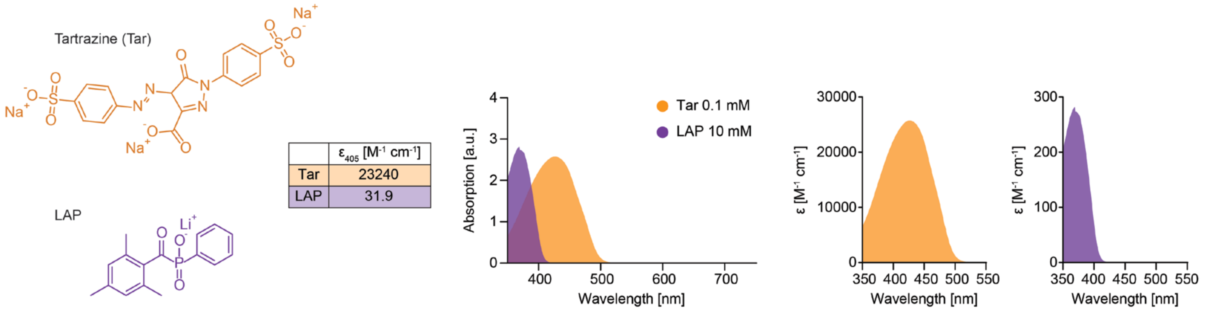


**Figure S7**. Molecular structures, absorption spectra. and extinction molar coefficients at 405 nm of photoinititor (LAP) and photoabsorber (tartrazine).

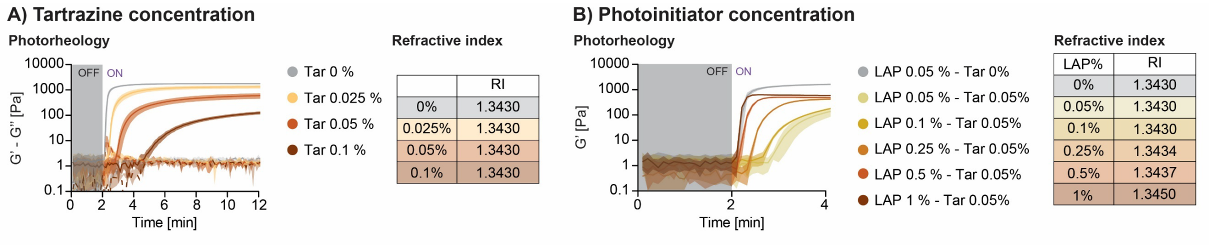


**Figure S8**. A) Photorheology analysis of the effect of tartrazine (Tar) concentration on 5% fGel-SH/NB, 0.05% LAP photoresin crosslinking kinetics and refractive index. B) Photorheology analysis of the effect of LAP concentration on 5% fGel-SH/NB photoresin crosslinking kinetics and refractive index.

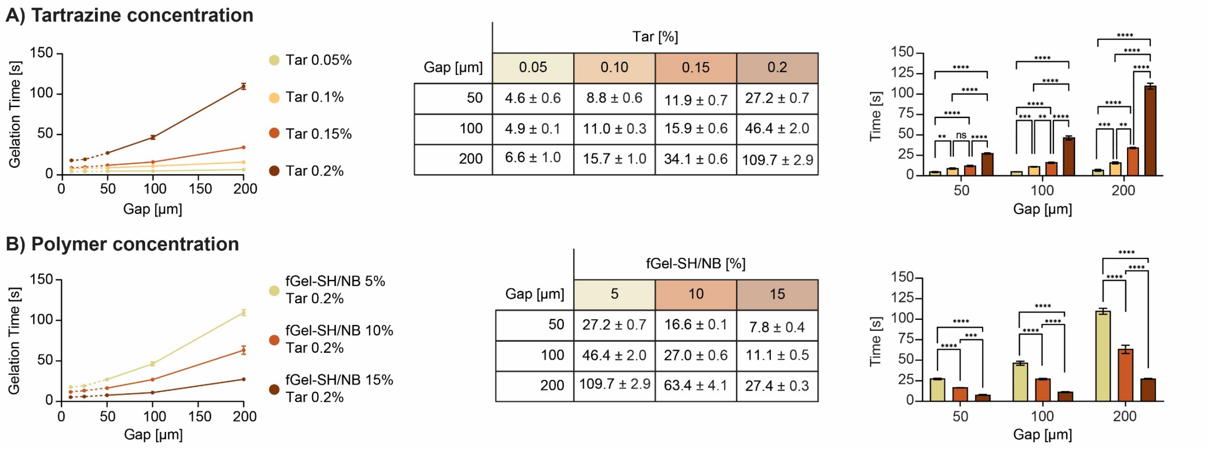


**Figure S9**. A) Gelation onset times for 5% fGel-SH/NB, 0.05% LAP photoresin at varying measurement gaps and photoabsorber (Tar) concentrations. B) Gelation onset times for fGel-SH/NB, 0.05% LAP, 0.2% Tar photoresin at varying measurement gaps and polymer concentrations.

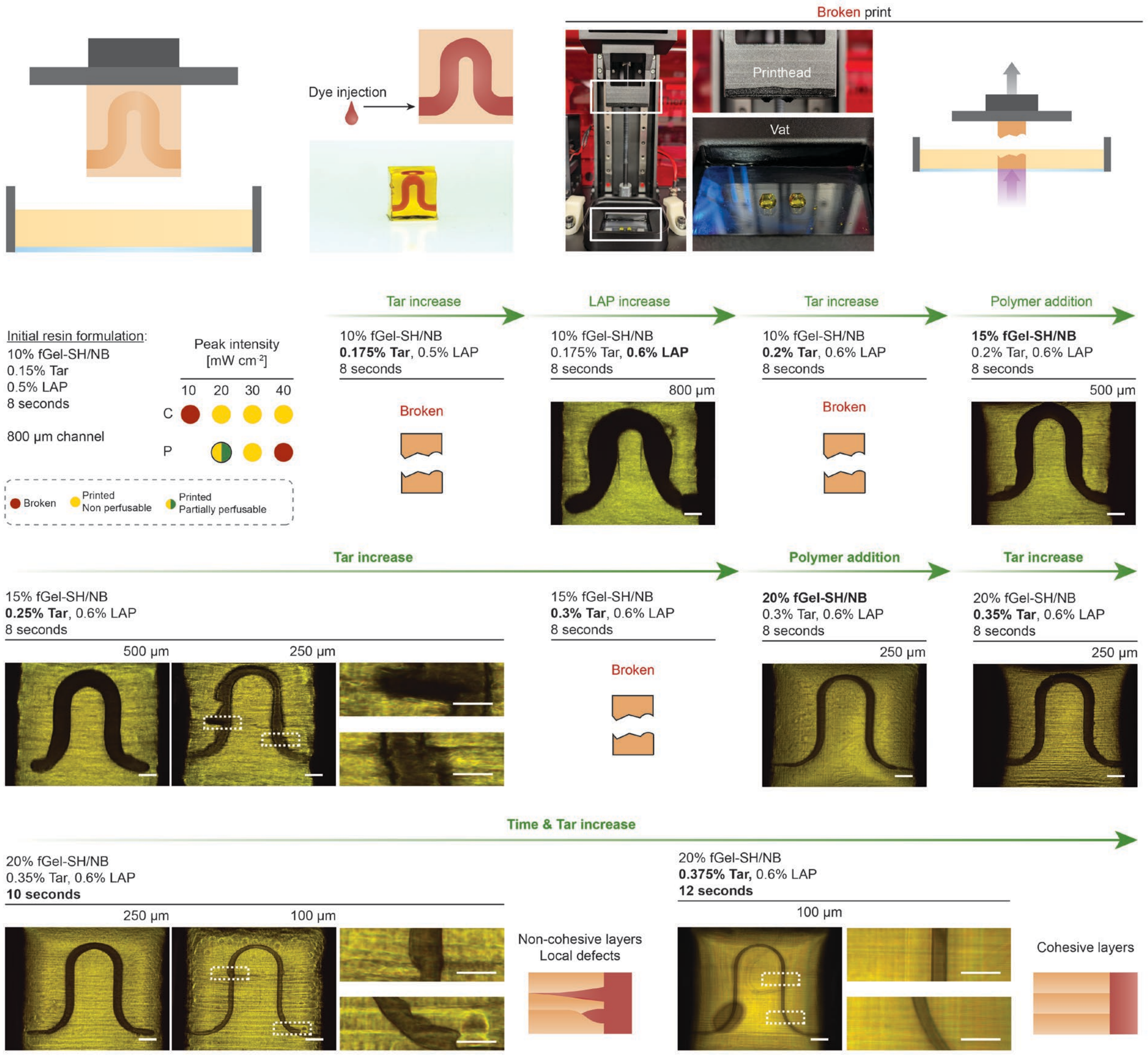


**Figure S10**. Photoresin and printing optimization steps to attain stable hydrogel constructs with an embedded microchannel (100 µm in diameter). Scale bars: 500 µm, close-ups: 250 µm.

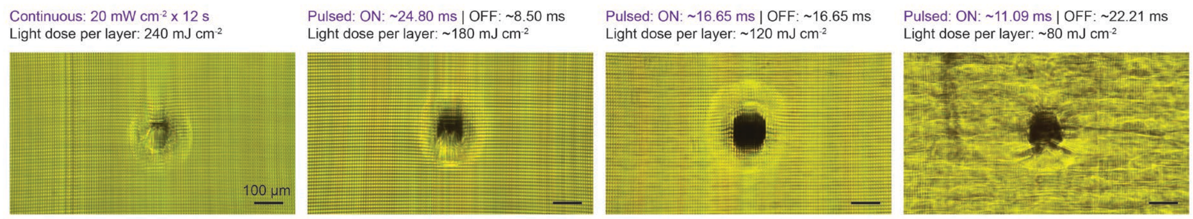


**Figure S11**. Aperture of horizontal microchannels (100 µm diameter) printed with continuous or pulsed illumination patterns.

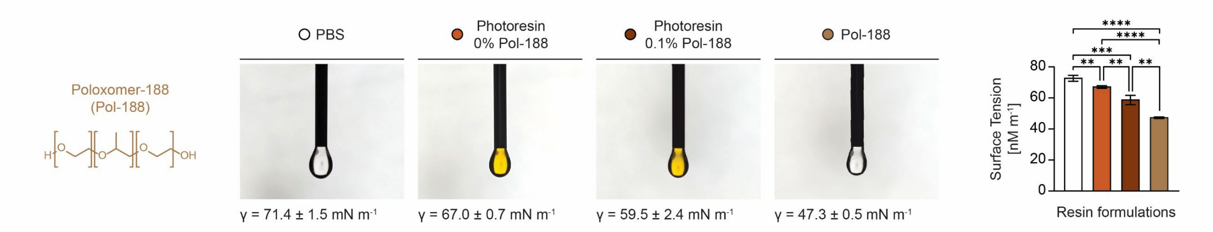


**Figure S12**. Pendant drop surface tension analysis of photoresin (20% fGel-SH/NB, 0.05% LAP, 0.375% Tar) without and with non-ionic surfactant Poloxamer-188 (Pol-188).

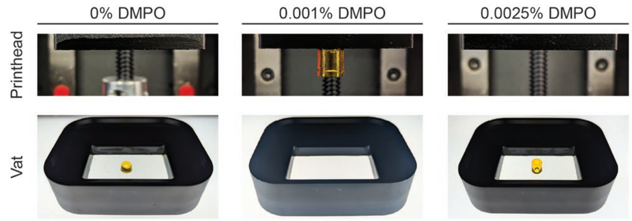


**Figure S13**. DLP printing of a 1 cm cylindrical construct (~ 5 h, 45 min) without and with DMPO at various concentrations.

**A) TEMPO**

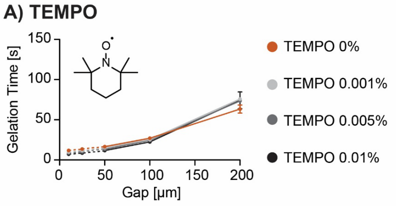


| Gap [µm] | TEMPO [%] 0 | 0.001 | 0.005 | 0.01 |
|---|---|---|---|---|
| 50 | 16.6 ± 0.16 | 15.3 ± 0.9 | 13.3 ± 1.1 | 11.7 ± 1.1 |
| 100 | 27.0 ± 0.6 | 25.4 ± 0.7 | 24.5 ± 0.9 | 22.4 ± 0.6 |
| 200 | 63.4 ± 4.1 | 75.9 ± 0.6 | 73.4 ± 1.0 | 74.2 ± 8.6 |

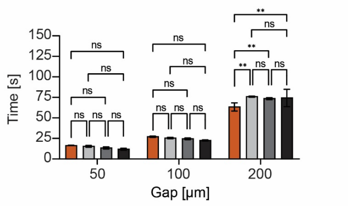


**B) DMPO**

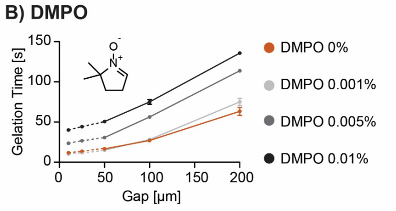


| Gap [µm] | DMPO [%] 0 | 0.001 | 0.005 | 0.01 |
|---|---|---|---|---|
| 50 | 16.6 ± 0.16 | 15.0 ± 0.2 | 30.7 ± 1.0 | 50.6 ± 0.9 |
| 100 | 27.0 ± 0.6 | 28.2 ± 0.6 | 56.3 ± 1.1 | 75.1 ± 2.4 |
| 200 | 63.4 ± 4.1 | 75.0 ± 3.6 | 113.8 ± 0.3 | 135.8 ± 0.6 |

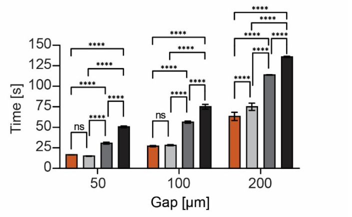


**Figure S14**. Gelation onset times for 10% fGel-SH/NB, 0.6% LAP photoresin at varying measurement gaps and TEMPO (A) or DMPO (B) concentrations.

| IDX % | RI |
|---|---|
| 0 % | 1.3654 |
| 10 % | 1.3809 |
| 20 % | 1.3908 |
| 30 % | 1.4054 |
| 60 %* | 1.4292 |

*undiluted OptiPrep™

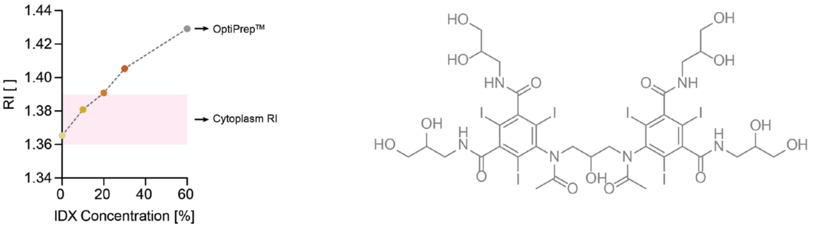


**Figure S15**. Tuning of the refractive index (RI) of the final resin formulation (20% fGel-SH/NB, 0.6% LAP, 0.375% Tar, 0.1% Pol-188, 0.001% DMPO) with addition of RI matching compound Iodixanol (IDX, structure on the right).

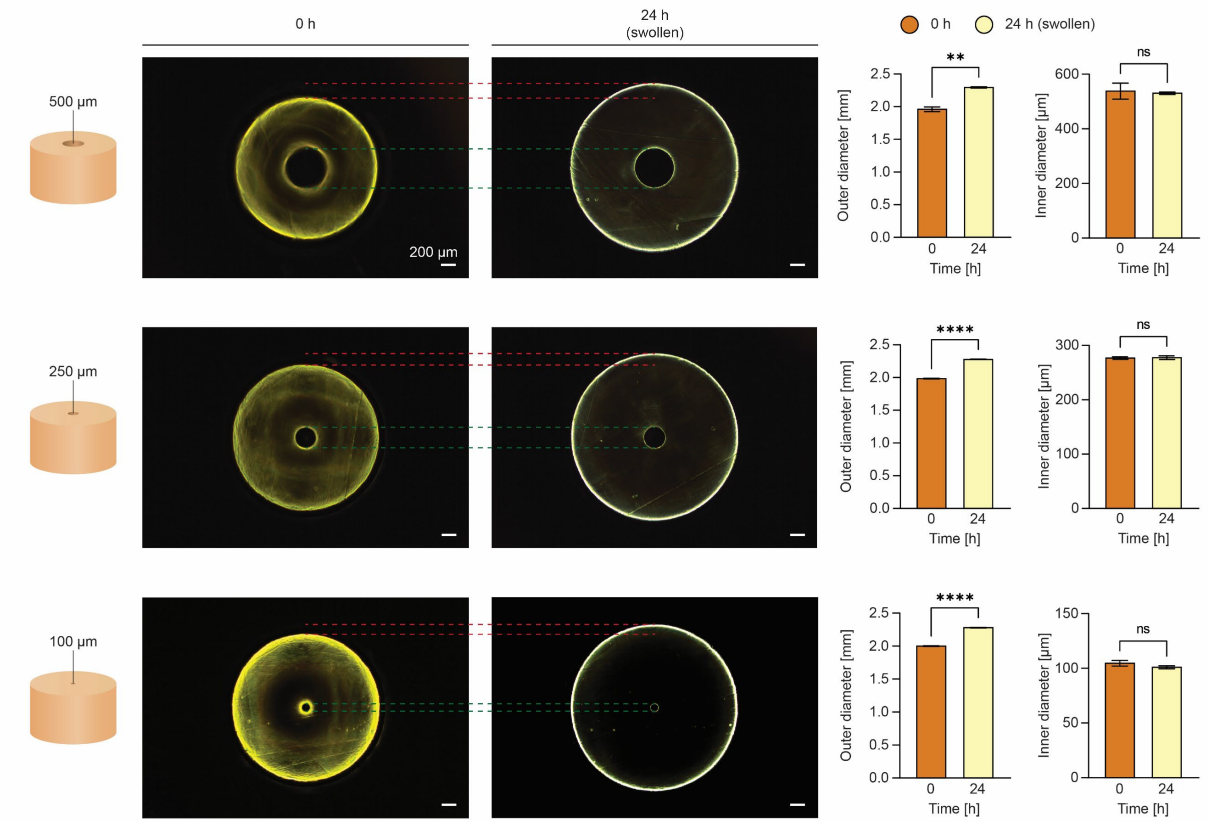


**Figure S16**. Swelling behavior of outer and inner diameter of printed hydrogel constructs with embedded microchannels of varying diameter.

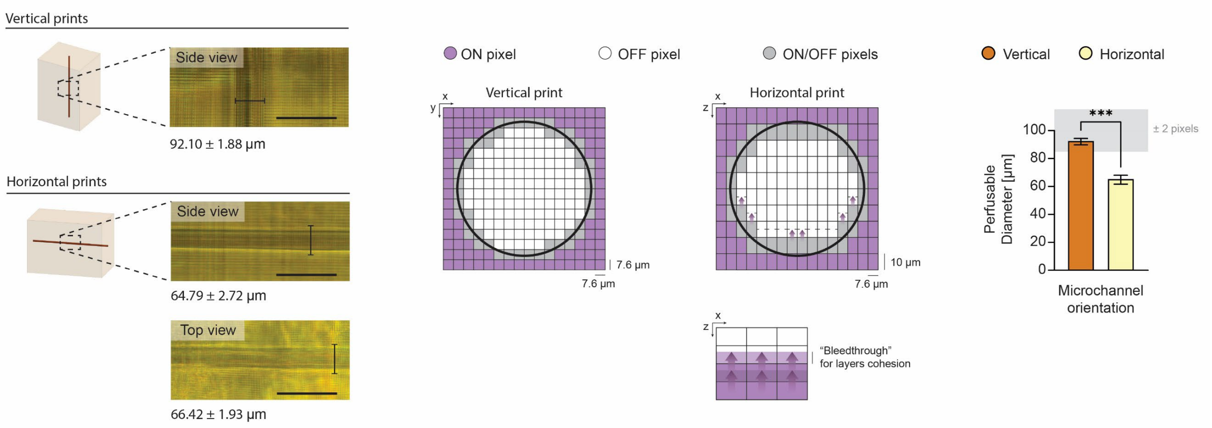


**Figure S17**. Details on mismatch between microchannel target diameter (100 µm) and experimental observation upon injection of filler. Scale bar: 200 µm.

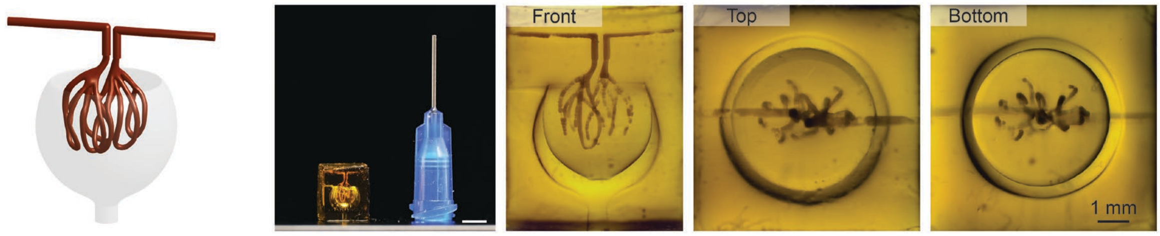


**Figure S18**. Schematic of biomimetic capillary network found in human kidney glomeruli and image of dye-perfused vasculature and air-filled Bowman's capsule. Scale bars: 4 mm (picture). On the right, microscopy close-ups of the printed construct.

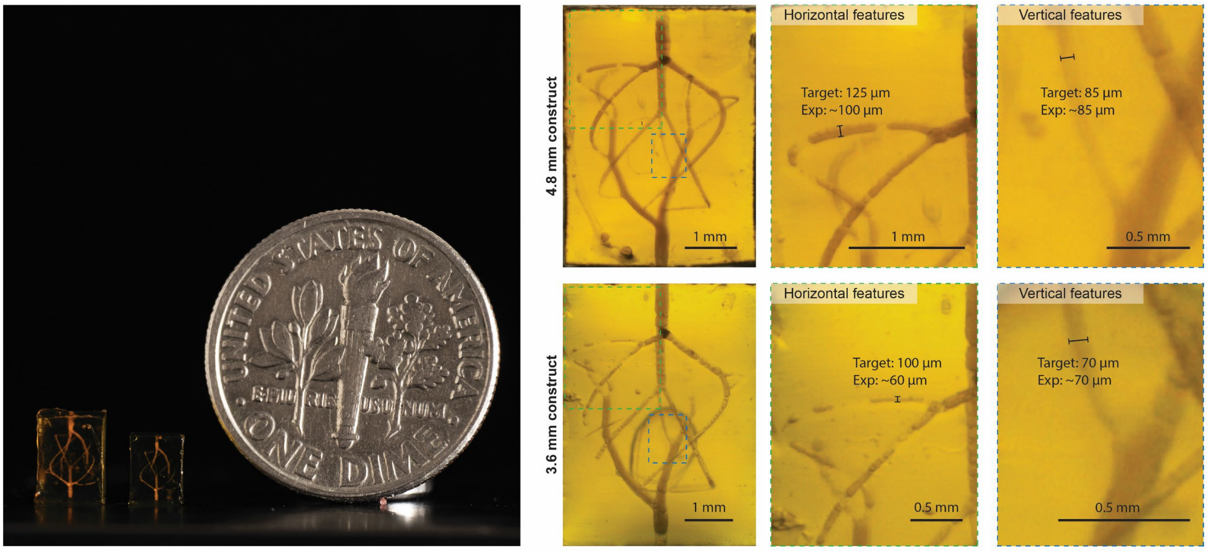


**Figure S19**. Biomimetic vascular network printed within hydrogel constructs of different size (left) (4.8 mm and 3.6 mm compared to the 6 mm print of Figure 4B) and microscopy close-ups (right).

**Table S1.** Parameters and resolutions of recent literature on DLP printing of perfusable horizontal channels in biocompatible hydrogels.

| Ref. | Photoresin | Photoinitiator | Photoabsorber | λ [nm] | Light Intensity [mW cm$^{-2}$] | Z-step [µm] | Pixel size [µm] | Min horizontal channel diameter [µm] |
|---|---|---|---|---|---|---|---|---|
| [21] | 10% PVA-MA<br>1% Gel-MA (DS 60%) | 0.2 mM Ru<br>2 mM SPS | 1% Ponceau4R | Vis | 7.25 | 50 | 30 | 500 |
| [8] | 20% PEG-DA (6 kDa) | 1% LAP | 0.05-0.5% Tartrazine | 405 | 15-20 | 25-200 | 10-50 | 350 |
| [20] | 10% PEG-DA (700 Da) | 0.1% LAP | 0.043% Chlorophyllin | 405 | / | 25-100 | 140 | 600 |
| [26] | 10% fGel-MA (DS 90%) | 2 mM Ru<br>20 mM SPS | 0.07% Ponceau4R | Vis | 6.5 | 50 | 30 | 180 |
| [43] | 8% PEG-DA (10 kDa) | 0.2% LAP | 0.5% R1800 | 365 | 6 | 50 | 30-65 | 750 |
| [22] | 5% Gel-MA (DS 81%)<br>3% HA-MA (100 kDa) | 1 mM Ru<br>10 mM SPS | 2% Ponceau4R | Vis | / | 100 | 50 | 1000 |
| [9] | 5% Gel-MA | 0.6% LAP | 1% yellow food dye | 405 | / | / | / | 250 |
| [34] | 0.6% HA-MA<br>4.5% HA-NB<br>0.25 mM Ad-CD | 0.5% LAP | 0.05% Tartrazine | 405 | 15 | 100 | 35 | 1000 |
| [35] | 3% Gel-NB<br>0.5% PEG4-NB (20 kDa)<br>5% PEG4-SH (10 kDa) | 0.29% LAP | 0.064% Tartrazine | 405 | 24.3 | 100 | 35 | 300 |
| [36] | 5% PEG8-NB (20 kDa)<br>4% PEG4-SH (10 kDa) | 0.29% LAP | 0.075% Tartrazine | 405 | 13.4-19.2 | 50-100 | 35 | 800-1000 |
| [44] | 60% Acrylamide<br>0.1% Bisacrylamide | 0.25% LAP | 0.075% Tartrazine | 405 | 20 | 100 | 35 | 600 |
| This work | 20% fGel-SH/NB | 0.6% LAP | 0.375% Tartrazine | 405 | 20 | 10 | 7.6 | 20 |
